# The Human Rectus Femoris Muscle Receives an Independent Common Synaptic Input During Isometric Leg Extensions

**DOI:** 10.64898/2026.07.14.738405

**Authors:** Finja Beermann, Daniel Haller, Luca Hofbeck, Simone Zaccaron, Marcel Betsch, Alessandro Del Vecchio

## Abstract

The distribution of common synaptic inputs across spinal motor neuron pools within and across synergist muscles reveals direct insight in the neural control of movement. We previously documented two dominant common inputs in the human vastus lateralis (VL) and medialis (VM) muscle during isometric leg extensions. Whether these inputs are also shared with rectus femoris (RF), a synergist muscle that contributes substantially to leg extension but is biarticular, remains unknown. We simultaneously recorded motor unit activity from VM, RF, and VL using multiple targeted intramuscular electromyographic sensors during isometric knee extension. Decomposed motor unit spike trains were analyzed using discharge characteristics, pairwise correlation, spectral coherence, explained variance, and factor analysis to characterize the structure of common synaptic input within and across muscles. Despite their shared mechanical output via a common patellar tendon, the three muscles exhibited distinct patterns of neural organization. RF consistently received a strong, low-dimensional common input that was largely independent of the drive to VM and VL, as evidenced by elevated within-muscle correlation and coherence, low between-muscle coupling, and an invariant dedicated latent factor across all participants. VM and VL shared a substantial proportion of common synaptic input, but the degree of coupling and the underlying factor structure varied across individuals, ranging from a single shared factor to largely independent muscle-specific inputs. These findings indicate that the quadriceps are organized in a muscle- and subject-specific manner, with RF receiving a dedicated independent input.

## INTRODUCTION

Coordinated movement requires the precise distribution of neural commands across multiple muscles. Motor neurons (MN) with their direct connection to muscle fibers serve as the final interface to convert these neural signals into muscle contraction (Liddell and Sherrington, 1925). They receive synaptic input from multiple sources, including afferent feedback, spinal interneurons, and descending supraspinal pathways (Gandevia, 2001; Heckman and Enoka, 2012). Using high-density surface electromyography (HDsEMG) or intramuscular electromyography (iEMG), the activity of single motor units (MUs) can be recorded directly, allowing the examination of common input architecture within and across muscles (Holobar and Zazula, 2007; Negro et al., 2009; Farina et al., 2014b,a; Negro et al., 2016a). Within individual muscles, strong common synaptic input has been consistently reported. For example, coherence analysis has demonstrated common input in the first dorsal interosseous (FDI) and tibialis anterior (TA) (Negro et al., 2016b; Dideriksen et al., 2018; Cabral et al., 2024), whereas correlation and principal component analyses have revealed similar findings for the gastrocnemius lateralis (GL) and gastrocnemius medialis (GM) (Levine et al., 2023). In the TA, individuals were unable to volitionally control single MUs independently, suggesting that MU activity is strongly constrained by common synaptic input (Bräcklein et al., 2022; Lee et al., 2026). Between muscles, the degree of shared input appears to depend on functional organization. Coherence between the GM and GL is minimal (Hug et al., 2021; Levine et al., 2023), and participants demonstrated a notable ability to dissociate their activity under online feedback (Rossato et al., 2024), indicating largely independent neural control. In contrast, the neural organization of the vastus medialis (VM) and vastus lateralis (VL) is less consistent and remains debated. Although these synergistic knee extensors exhibit high intermuscular coherence (Laine et al., 2015; Avrillon et al., 2021) and could not be dissociated under online feedback using HDsEMG decomposition (Rossato et al., 2024), Haller et al. (2026) demonstrated that targeted intramuscular EMG feedback enabled voluntary separation in 9 out of 10 participants. Additionally, Dernoncourt et al. (2025) found that the VL MU pool alone was best described by three distinct latent factors, indicating considerable variability in the inferred neural organization depending on the analytical approach.

Within the quadriceps, only limited data exist regarding the neural organization of the biarticular RF muscle. One of the few studies simultaneously examining VM, RF, and VL reported preliminary evidence of an independent neural drive to RF (Rossato et al., 2022), though the small number of identified MUs constrained inference at the pool level. Nevertheless, this interpretation aligns with earlier work demonstrating independent control of quadriceps heads, where experimentally induced pain in RF selectively reduced RF EMG activity without affecting VM during isolated knee extension (Hug et al., 2014). Importantly, previous studies largely relied on surface EMG, which is prone to cross-talk, limited in detecting deep or small MUs, and captures only a localized region of the muscle, thereby not capturing its full spatial extent (Germer et al., 2021). These recordings can reveal muscle-level synergies but cannot determine how inputs are distributed across individual MUs. The first study to investigate the number of latent factors expressed in motor unit modes (Del Vecchio et al., 2023) found that a small proportion of MUs deviated from the dominant muscle-specific input. More recently, Weinman et al. (2026) identified distinct MU modes in the proximal and distal regions of the TA using two HDsEMG grids, and Haller et al. (2026) similarly reported that distal VM activity more closely resembled VL than proximal VM using multiple intramuscular implants. These limitations highlight the need to assess MU-level input organization spanning proximal to distal regions across all three quadriceps heads.

To address this question, we simultaneously recorded MU activity from VM, RF, and VL using multiple targeted iEMG electrodes during isometric knee extension. This task emphasizes their shared function as knee extensors while minimizing the influence of RF’s biarticular role at the hip. Discharge characteristics, pairwise correlation, spectral coherence, explained variance, and factor analysis were used to characterize the organization of common synaptic input within and across the quadriceps muscles. We hypothesized that RF would exhibit a distinct pattern of common synaptic input relative to VM and VL, reflecting an independent neural drive even during a task that promotes synergistic knee extension. Furthermore, given the debated organization of VM and VL, we expected that including RF as a functionally distinct muscle would help to better characterize the organization of shared input within the quadriceps.

## MATERIAL AND METHODS

### Participants and ethical approval

A total of ten healthy participants (28.6 ± 4.6 years [*µ* ± SD], 7 males, 3 females) volunteered for the study. None reported any known neurological, neuromuscular, or musculoskeletal disorders. All participants gave their informed written consent before participating. The study was approved by the local Ethics Committee of the Friedrich-Alexander University Erlangen-Nürnberg (approval no. 25-37 2-S) according to the Declaration of Helsinki.

### Signal Recordings

Intramuscular EMG activity was recorded from VM, RF, and VL of the participant’s self-reported dominant leg. Under ultrasound guidance by an experienced physician, the proximal and distal ends of each muscle were identified. Three bipolar hook-wire electrodes (Stainless Steel/Ag; diameter 0.11 mm; SpesMedica, Battipaglia, Italy) were inserted into each muscle and distributed across the proximal, central, and distal regions to a depth of approximately 2 cm. An ankle strap reference (Bioelettronica, Turin, Italy) was placed around the ankle of the dominant leg. A customized knee dynamometer with an attached three-axis force sensor (K3D120, 1 kN, accuracy class 0.5; ME-Meßsysteme, Hennigsdorf, Germany) was used to record the applied force of knee extension. The participants were seated upright with the hip flexed to 90°and the knee positioned at 45°angle flexion (0°defined as full knee extension). The lower leg was secured to the force sensor with two adjustable straps placed over the shin. The position of the sensor with respect to the leg was adjusted for each participant to around 10 cm above the ankle. All data were recorded with a sampling frequency of 10,240 Hz using the acquisition system Quattrocento (OT Bioelettronica, Turin, Italy). EMG signals were subsequently filtered with a second-order Butterworth bandpass filter between 100 and 4,400 Hz.

### Experimental Protocol

Before needle implantation, participants performed three measurements of maximal voluntary contraction (MVC), with 1 min of rest between trials. Participants were instructed to ‘push as hard as possible’ for 3 to 5 s, while the maximum force of the previous trial was displayed on a monitor. In addition, the participants received strong verbal encouragement. The greatest force recorded during any of the three MVCs was used as a reference for visual force feedback of submaximal contractions. MVC measurements were performed prior to needle implantation, as discomfort or pain associated with the intramuscular electrodes could potentially influence maximal force production. Subsequently, participants performed a series of isometric ramp contractions. Each ramp consisted of a linear force increase to the target level at 5% MVC *s*^*−*1^, a constant-force plateau of 40, 30, or 20 s at 5%, 15%, or 30% MVC, respectively, and a linear decrease at the same rate. The order of submaximal force levels was randomized and separated by 2 min to avoid fatigue. Although contractions were recorded at all three force levels, only the 30% MVC condition was included in the subsequent analysis, as at lower force levels the number of decomposed MUs in the RF was insufficient (*<*4) in the majority of participants. Additionally, two participants were excluded from the analysis due to an insufficient number of decomposed MUs in the RF at 30% MVC, resulting in a final sample of 8 participants (5 males, 3 females).

### Motor Unit Decomposition

Intramuscular EMG signals were decomposed into individual discharge times of MU using the open-source decomposition software EMGLAB (McGill et al., 2005), running in MATLAB (MATLAB R2022b; MathWorks Inc., Natick, MA). Prior to decomposition, signals were high-pass filtered at a cut-off frequency of 1000 Hz to remove low-frequency noise. Each of the nine iEMG channels was decomposed individually. The algorithm initially detects candidate spikes and clusters them based on waveform similarity using principal component analysis (PCA) and k-means clustering to form initial templates. These templates are then iteratively refined through template matching and supervised manual verification to account for overlapping potentials and waveform variability. All detected MUs were visually inspected, and missed discharges or superpositions were manually corrected where necessary. MUs exhibiting an average action potential amplitude less than twice the baseline noise level were excluded to ensure that only reliably detected MUs were retained for further analysis.

### Data Analysis

All analyses were restricted to the plateau phase (excluding the first and last 3 s), with inactive MUs or those with a coefficient of variation (CoV) of the interspike interval greater than 2 excluded to ensure firing stability. The remaining spike trains were used to calculate the mean discharge rate (DR) and the CoV of each MU. To characterize the degree to which MU populations of the three muscles are distinguishable in terms of their neural drive, a complementary set of analyses was applied, including discharge characteristics, pairwise correlation, spectral coherence, explained variance, and factor analysis. Additionally, t-distributed Stochastic Neighbor Embedding (t-SNE) was applied to visualize the multivariate separation of MU discharge patterns across muscles in a low-dimensional projection.

#### Motor Unit Correlation

To quantify the strength of low-frequency common drive (De Luca and Erim, 1994), MU binary spike trains were convolved with a 400 ms Hanning window (no phase shift), corresponding to a 2.5 Hz low-pass filter (De Luca et al., 1982; Myers et al., 2004), and subsequently z-scored. For each unique MU pair within (VM, RF, VL) and between muscles (VM–RF, VM–VL, RF–VL), the Pearson’s correlation coefficient was calculated (Del Vecchio et al., 2023). Given that all muscles are innervated by the femoral nerve, inter-muscular propagation delays are negligible relative to the 400 ms smoothing kernel. To verify robustness, analyses were repeated on non-normalized signals and using cross-correlation within a ± 100 ms window (Hug et al., 2023), yielding only minor differences that did not alter any statistically significant findings.

#### Motor Unit Pool Coherence

Coherence spectra were calculated for MU pools within (VM, RF, VL) and across muscles (VM–RF, VM–VL, RF–VL) using the Neurospec 2.11 toolbox (www.neurospec.org; Halliday, 2015) implemented in MATLAB. MUs were randomly divided into two equal subpools, and binary spike trains within each subpool were summed to obtain cumulative spike trains (CSTs), which were linearly detrended prior to analysis. Spectral estimates were computed using a multi-taper approach (time–bandwidth product of 2; frequency resolution of 0.625 Hz). This procedure was repeated 100 times with random MU permutations, and coherence spectra were averaged across iterations (Hug et al., 2021; Negro et al., 2016b). To correct for bias, the mean coherence within the 250–500 Hz range, where no physiologically meaningful coherence is expected, was subtracted from all frequency bins, and resulting negative values were set to zero (Castronovo et al., 2015; Maillet et al., 2022; Cabral et al., 2024). Following bias correction, the area under the curve (AUC) and peak coherence were computed for the delta (0–5 Hz) and alpha (5–13 Hz) bands. AUC values were normalized to the respective bandwidth, yielding coherence estimates in units of Hz^-1^ to allow direct comparison between frequency bands. Because coherence depends on the number of MUs included in the analysis (Negro and Farina, 2012; Farina et al., 2014b; Dideriksen et al., 2018; Castronovo et al., 2015), coherence was calculated for increasing pool sizes using identical sizes for within- and between-muscle comparisons. As a third measure, the proportion of common synaptic input (PCI) (Negro et al., 2016b) was estimated by modeling peak coherence as a function of pool size using nonlinear least-squares optimization. The PCI was defined as 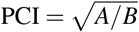 where A and B denote the power of the common and independent synaptic input, respectively, with higher values indicating a greater proportion of shared input. For full derivation see Negro et al. (2016b).

#### Cumulative Explained Variance

Complementary to the correlation and coherence analyses described above, cumulative explained variance was computed to compare the dimensionality of MU discharge activity across muscles, using both principal component analysis (PCA; Levine et al. (2023)) and factor analysis (FA; Del Vecchio et al. (2023); Dernoncourt et al. (2025); Weinman et al. (2024)). PCA was performed via eigendecomposition of the correlation matrix of the z-scored, Hanning-convolved spike trains, providing a full decomposition of total variance. FA was performed using maximum likelihood estimation with promax rotation (Dernoncourt et al., 2025), allowing the extracted factors to be correlated rather than enforcing orthogonality, consistent with the shared innervation and mechanical coupling of the three muscles. For both approaches, cumulative explained variance was computed retaining the first eight components or factors, and to control for differences in the number of MUs across muscles, 100 random subsets of size equal to the minimum MU count across muscles were drawn per participant, with the resulting curves averaged across repetitions. To verify robustness, the FA was repeated using orthogonal varimax rotation, yielding only minor differences that did not alter any statistically significant findings.

#### Low-Dimensional Projection

t-distributed Stochastic Neighbor Embedding (t-SNE) is a nonlinear dimensionality reduction technique that projects high-dimensional data into a low-dimensional space while preserving local neighborhood structure (van der Maaten and Hinton, 2008). It was applied separately for each participant to the z-scored, Hanning-convolved spike trains to visualize the multivariate separation of MU discharge patterns across muscles. Perplexity was set to min(30, ⌊*n/*3⌋ *−*1), where *n* is the number of available MUs per participant, with a minimum of 2. A fixed random seed (42) ensured reproducibility. As inter-point distances in the projected space are not metrically interpretable, the resulting projections served as a visual complement to the correlation and coherence analyses only, and no formal statistical analysis was performed.

#### Factor Analysis

Del Vecchio et al. (2023) showed that MUs can form functional groups spanning multiple muscles, indicating the presence of MU modes that reflect shared neural drive across muscles. To investigate these MU modes across all three muscles, FA was applied to the pooled, z-scored spike trains. Factor extraction was performed using maximum likelihood estimation with promax rotation. Unlike previous studies that applied a fixed factor number, the optimal number of factors was determined individually per participant using a sequential three-step procedure, with a maximum of nine candidate factors based on Dernoncourt et al. (2025), who reported up to three latent factors within a single muscle. First, parallel analysis (Horn, 1965), implemented using the permutation-based variant proposed by Buja and Eyuboglu (1992), retained only factors with eigenvalues exceeding the 95th percentile of a null distribution generated from 200 column-permuted datasets. Second, only factors explaining at least 5% additional variance beyond the previous model were retained (Clark et al., 2010). Third, factors were validated against a surrogate threshold derived from 100 datasets with independently circularly-shifted spike trains, preserving interspike interval statistics while eliminating inter-MU covariance. A factor was accepted only if its explained variance exceeded the 95th percentile of this distribution. The final factor number was the largest satisfying all three criteria. Following factor extraction, each factor was assigned a dominant muscle based on the highest mean absolute loading across MUs of that muscle. If fewer than three factors were extracted, a single factor may span multiple muscles, if more than three were extracted, a single muscle may be represented by multiple factors. Each MU was then assigned to the factor on which it showed the highest absolute loading. For each muscle pool, the percentage of MUs assigned to each muscle-specific factor (VM–factor, RF–factor, VL–factor) was computed, reflecting the degree to which each muscle’s MUs are driven by a common input shared within or across muscles.

## Statistical Analysis

All statistical analyses were performed in MATLAB. Mean DR, CoV, Pearson’s correlation coefficients, and cumulative explained variance (PCA and FA) were compared across muscles (VM, RF, VL) and muscle-pairs (VM–RF, VM–VL, RF–VL) using linear mixed-effects models (LME), with muscle or muscle-pair as a fixed effect and subject-by-muscle interaction as random intercepts. For cumulative explained variance, separate models were fitted for PCA- and FA-derived estimates, each additionally including number of components or factors and its interaction with muscle as fixed effects. Pearson’s correlation coefficients were Fisher z-transformed prior to statistical analysis to stabilize variance and approximate a normal distribution, as raw correlation values are bounded and asymmetrically distributed. As cumulative explained variance represents a bounded proportion rather than a correlation-based measure, values were logit-transformed prior to analysis, with boundary values clipped to [0.1%, 99.9%]. The significance of fixed effects for all LME was assessed using ANOVA on the fitted LME models, with effect sizes reported as *Cohen*^*′*^*s f* ^2^ (small: *f* ^2^ ≥ 0.02, medium: *f* ^2^ ≥ 0.15, large: *f* ^2^ ≥ 0.35). Post-hoc pairwise comparisons were conducted using coefficient tests and effect sizes are reported as Cohen’s d (small: *d* ≥ 0.2, medium: *d* ≥ 0.5, large: *d* ≥ 0.8). As the large number of MU pairs in the correlation analysis can result in statistically significant but practically negligible effects, post-hoc results for correlations were only reported when effects exceeded a predefined minimum Cohen’s d threshold, defined as *d >* 0.5. For cumulative explained variance, Holm correction was applied separately for PCA and FA across all 24 post-hoc comparisons (8 component or factor levels ×3 muscle pairs) simultaneously. For coherence-based measures, AUC and peak in the delta and alpha bands were extracted at matched MU pool sizes to avoid confounding effects of pool size and Fisher Z-transformed prior to analysis. As PCI is a model-derived estimate of shared input and its values are not constrained to the [-1, 1] interval, they were not Fisher Z-transformed. Differences between muscles (VM, RF, VL) and between muscle-pairs (VM–RF, VM–VL, RF–VL) were analyzed separately and assessed using Friedman tests, with effect sizes quantified via Kendall’s *W* (small: *W* ≥ 0.1, medium: *W* ≥ 0.3, large: *W* ≥ 0.5). Post-hoc pairwise comparisons were performed using Wilcoxon signed-rank tests with effect sizes reported as rank-biserial correlation *r*_*rb*_. Statistical significance was set at *α* = 0.05, with p-values adjusted using the Holm–Bonferroni procedure for multiple comparisons.

## RESULTS

During isometric knee extension at 30% MVC, MU activity was simultaneously recorded from VM, RF and VL using targeted intramuscular implants. A complementary set of analyses including discharge characteristics, pairwise correlation, spectral coherence, explained variance, and factor analysis was applied to the decomposed MU pools to characterize the neural input organization within (VM, RF, VL) and across (VM–RF, VM–VL, RF–VL) the quadriceps muscles. A total of 323 MUs (VM: n=133; RF: n=74; VL: n=116) were successfully decomposed, with on average 16.6 *±* 5.4, 9.2 *±* 5.6, and 14.5 *±* 6.5 MUs per participant for VM, RF, and VL, respectively.

### Discharge Properties

Mean DR and CoV were compared across muscles as a basic characterization of MU firing behavior. Mean DR differed significantly across muscles, *F*(2, 320) = 48.8, *p <* 0.001, Cohen’s *f* ^2^ = 0.44. The mean DR was 10.2 ± 1.6, 13.2 ± 1.7, and 9.3 ± 1.3 Hz for VM, RF, and VL, respectively (Figure 2 B, left panel). Post-hoc comparisons indicated that RF exhibited a significantly higher DR than both VM (difference = 2.95 Hz, *p <* 0.001), and VL (difference = 3.80 Hz, *p <* 0.001). VM and VL also differed significantly, with VM showing a higher DR than VL (difference = 0.85 Hz, *p* = 0.020). Discharge variability, as indexed by CoV, likewise differed significantly across muscles, *F*(2, 320) = 21.1, *p <* 0.001, Cohen’s *f* ^2^ = 0.23. Mean CoV was 10.1% ± 2.0%, 15.3% ± 3.8%, and 10.3% ± 2.1% for VM, RF, and VL, respectively (Figure 2 B, right panel). Post-hoc comparisons indicated a higher CoV in RF compared to both VM (difference = 4.6%, *p <* 0.001), and VL (difference = 4.5%, *p <* 0.001), whereas VM and VL did not differ significantly (*p* = 0.841).

**Figure 1.**
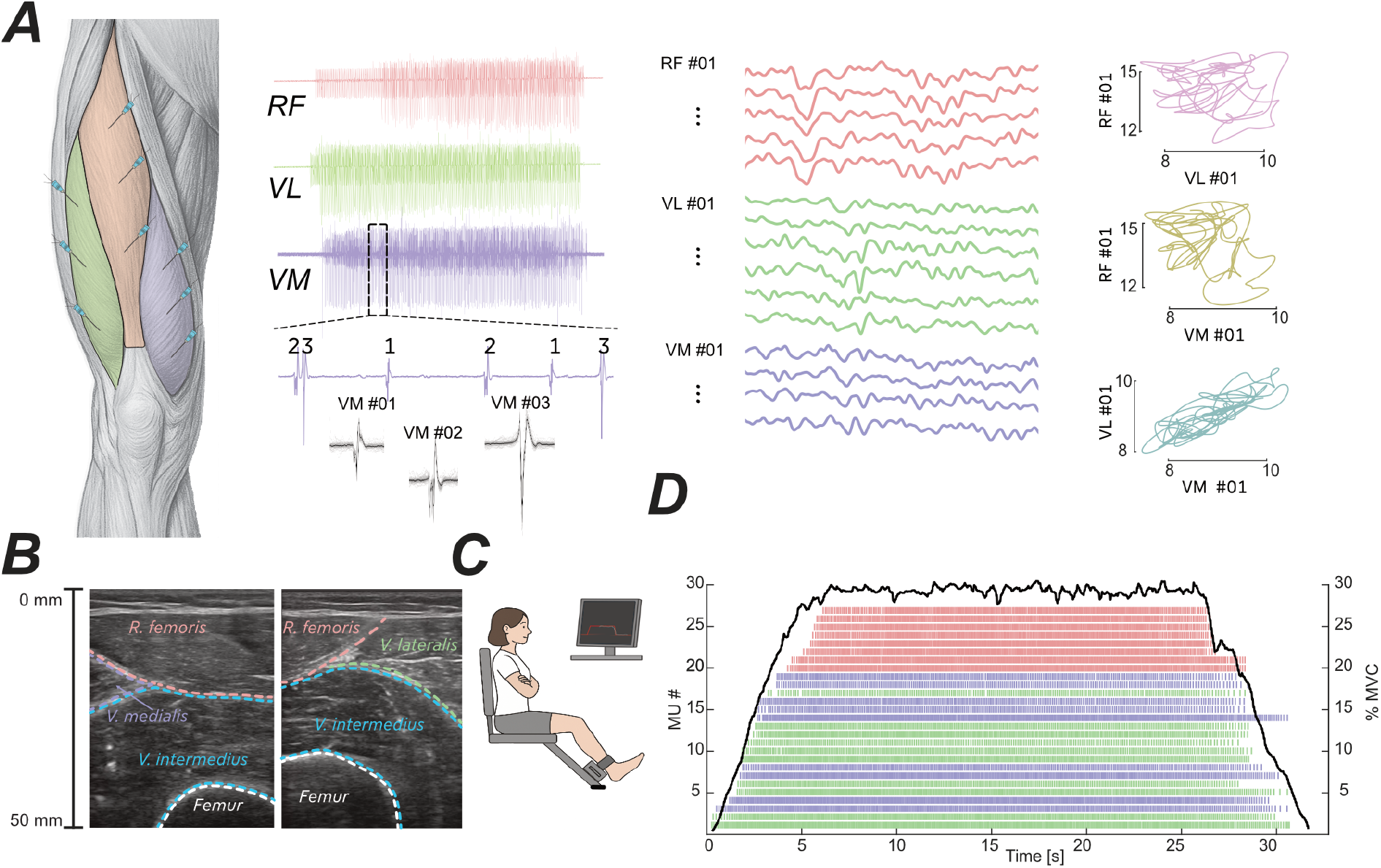
Three intramuscular bipolar fine-wire electrodes were inserted into the vastus medialis (VM), rectus femoris (RF), and vastus lateralis (VL) of the quadriceps (A, left panel), with electrode placement guided by ultrasound imaging (B). Participants performed isometric knee extensions at 30% MVC in a custom setup (C). Raw intramuscular EMG signals were decomposed into individual motor unit discharge times. Binary spike trains were convolved with a 400 ms Hanning window to derive smoothed discharge rate trajectories (A, middle panel). Biplots of smoothed discharge rates from MU pairs across muscles illustrate that some MUs co-vary closely, whereas others show little co-variation, suggesting independent neural drive (A, right panel). MU spike trains from an exemplary participant are shown sorted by recruitment order with the force trace depicted in black above (D). Color coding corresponds to muscle identity throughout: VM (purple), RF (pink), VL (green).

**Figure 2.**
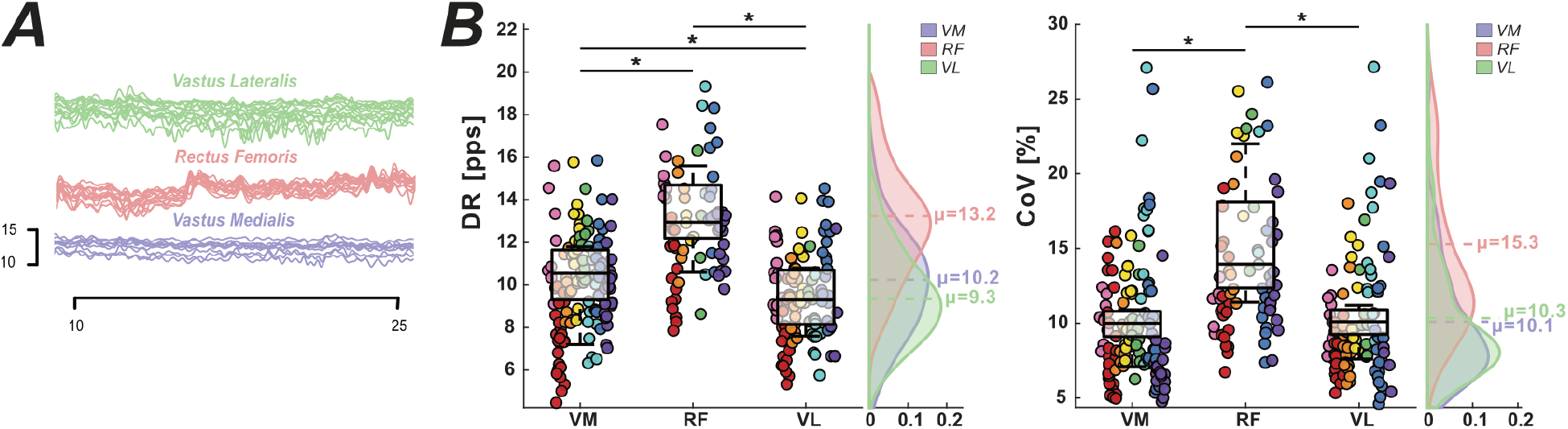
Smoothed discharge rate trajectories from all recorded MUs per muscle are shown for an exemplary participant (A). Mean discharge rate (DR; left panel) and coefficient of variation of inter-spike intervals (CoV; right panel) were calculated for each individual MU across all participants, displayed as scatter plots with overlaid box plots and a marginal density plot per muscle (B). Colors indicate muscle identity throughout: VM (purple), RF (pink), VL (green), with individual colors within each muscle representing different participants. Asterisks (*) denote statistically significant differences following Holm correction (*α* = 0.05).

### Motor Unit Correlations

To characterize the degree to which MU spike trains co-varied within and across muscles, Pearson’s correlation coefficients were computed for all unique MU pairs within (VM, RF, VL) and across muscle pairs (VM–RF, VM–VL, RF–VL). Within-muscle correlation strength differed significantly across muscles, *F*(2, 2485) = 559.1, *p <* 0.001, Cohen’s *f* ^2^ = 0.27. Post-hoc comparisons indicated higher correlations within RF than within VM (Δ*z* = 0.52, Cohen’s *d* = 1.78, *p <* 0.001) and within VL (Δ*z* = 0.57, *d* = 1.95, *p <* 0.001), whereas VM and VL did not differ significantly (*p* = 0.63). Between-muscle correlation strength also differed significantly across muscle pairs, *F*(2, 4475) = 937.1, *p <* 0.001, Cohen’s *f* ^2^ = 0.16. Post-hoc comparisons revealed higher correlations for VM–VL than for VM–RF (Δ*z* = 0.29, *d* = 1.17, *p <* 0.001) and RF–VL (Δ*z* = 0.36, *d* = 1.48, *p <* 0.001), while VM–RF and RF–VL did not differ significantly (*p* = 0.1). The distribution of correlation coefficients within and across muscles is illustrated in Figure 3 A and B. Beyond these systematic differences, substantial inter-individual variability was observed (Figure 3B). While muscle pair explained 13.4% of total variance (marginal *R*^2^), the full model accounted for 50% (conditional *R*^2^), with between-subject differences contributing the majority of explained variance (ICC = 0.52).

**Figure 3.**
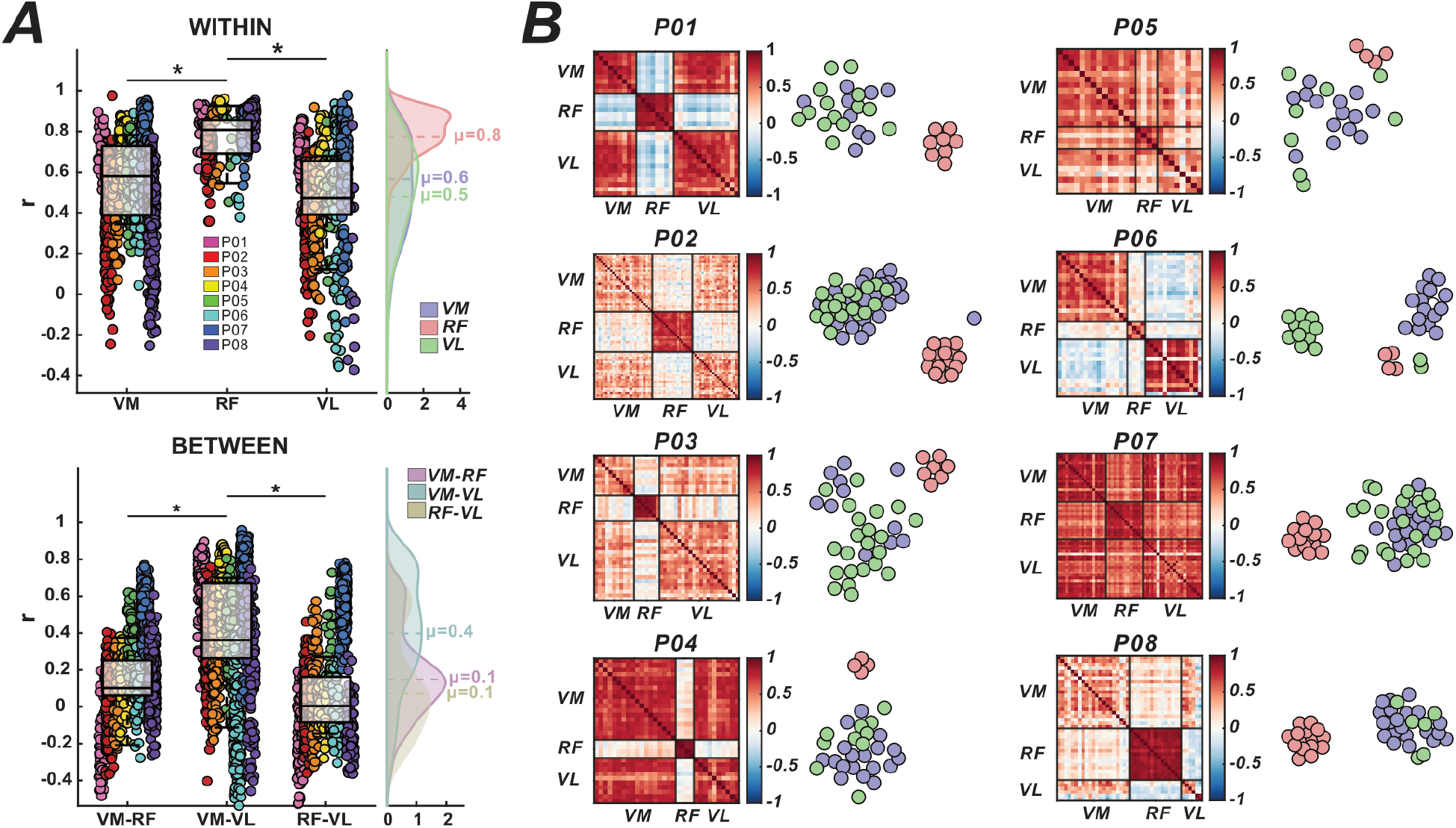
Binary spike trains were convolved with a 400 ms Hanning window, z-scored, and Pearson correlation coefficients computed for all unique MU pairs. Within-muscle pairwise correlation values are shown in the upper panel of (A) and across-muscle correlation values in the lower panel of (A), each displayed as scatter plots with overlaid box plots and a marginal density plot, aggregated across all participants. Colors indicate muscle identity throughout: VM (purple), RF (pink), VL (green). Asterisks (*) denote statistically significant differences following Holm correction (*α* = 0.05). For each participant (P01–P08), pairwise MU correlation matrices are displayed alongside t-SNE projections of the full MU pool, computed on z-scored spike trains, illustrating inter-individual differences in clustering structure (B).

### Low-Dimensional Visualization

To complement these analyses, t-SNE projections of the full MU pool are shown in Figure 3 B for each participant. The projections illustrate the multivariate separation of MU discharge patterns across muscles and are broadly consistent with the correlation structure. Across participants, RF formed a consistently separate cluster. The degree of overlap between VM and VL, in contrast, varied considerably between individuals, ranging from largely separate clusters (e.g., P 06) to substantial overlap (e.g., P 01).

### Motor Unit Pool Coherence

Complementing the time-domain correlation analysis, coherence quantifies shared neural drive in the frequency domain, enabling the identification of common oscillatory inputs within and across muscles. Coherence was predominantly concentrated in the delta band (0–5 Hz), as illustrated by the averaged spectra across participants (Figure 4 A). No significant differences were observed in the alpha band, subsequent analyses therefore focus only on the delta band. To also compare coherence across participants, normalized AUC coherence was computed across increasing pool sizes (Figure 4 B). The slope across participants varied considerably within each muscle and muscle pair, reflecting substantial inter-individual differences in common synaptic input. Summary coherence metrics (AUC, peak coherence, and PCI) for within- and across-muscle conditions are displayed in Figure 4 C, further illustrating the consistently elevated coherence in RF relative to VM and VL, and the stronger coupling between VM–VL compared to muscle pairs involving RF. Significant differences in coherence estimate were observed across muscles for all three measures, AUC (*χ*^2^(2) = 12.25, *p* = 0.0022, Kendall’s *W* = 0.77), peak coherence (*χ*^2^(2) = 9.25, *p* = 0.0098, Kendall’s *W* = 0.58), and PCI (*χ*^2^(2) = 9.25, *p* = 0.0098, Kendall’s *W* = 0.58). Post-hoc comparisons revealed a consistent pattern across all three measures. RF showed higher values than both VM (AUC: Δ*z* = 0.27, peak: Δ*z* = 0.42, PCI: Δ*z* = 0.53, all *p* ≤ 0.016) and VL (AUC: Δ*z* = 0.26, peak: Δ*z* = 0.41, PCI: Δ*z* = 0.51, all *p* ≤ 0.008), whereas VM and VL did not differ significantly across any measure (AUC: *p* = 0.74, peak: *p* = 0.95, PCI: *p* = 0.95). Coherence estimates across muscle pairs differed significantly for AUC (*χ*^2^(2) = 7.00, *p* = 0.0302, Kendall’s *W* = 0.44) and peak coherence (*χ*^2^(2) = 9.75, *p* = 0.0076, Kendall’s *W* = 0.61), but not for PCI (*χ*^2^(2) = 5.25, *p* = 0.0724, Kendall’s *W* = 0.33). Post-hoc comparisons for AUC and peak coherence revealed higher coherence for VM–VL compared to VM–RF (AUC: Δ*z* = 0.12, *p* ≤ 0.016, peak coherence: Δ*z* = 0.31, *p* ≤ 0.008), and compared to RF–VL (AUC: Δ*z* = 0.10, *p* ≤ 0.023, peak coherence: Δ*z* = 0.27, *p* ≤ 0.016), while VM–RF and RF–VL did not differ significantly (AUC: *p* = 0.38, peak coherence: *p* = 0.05).

**Figure 4.**
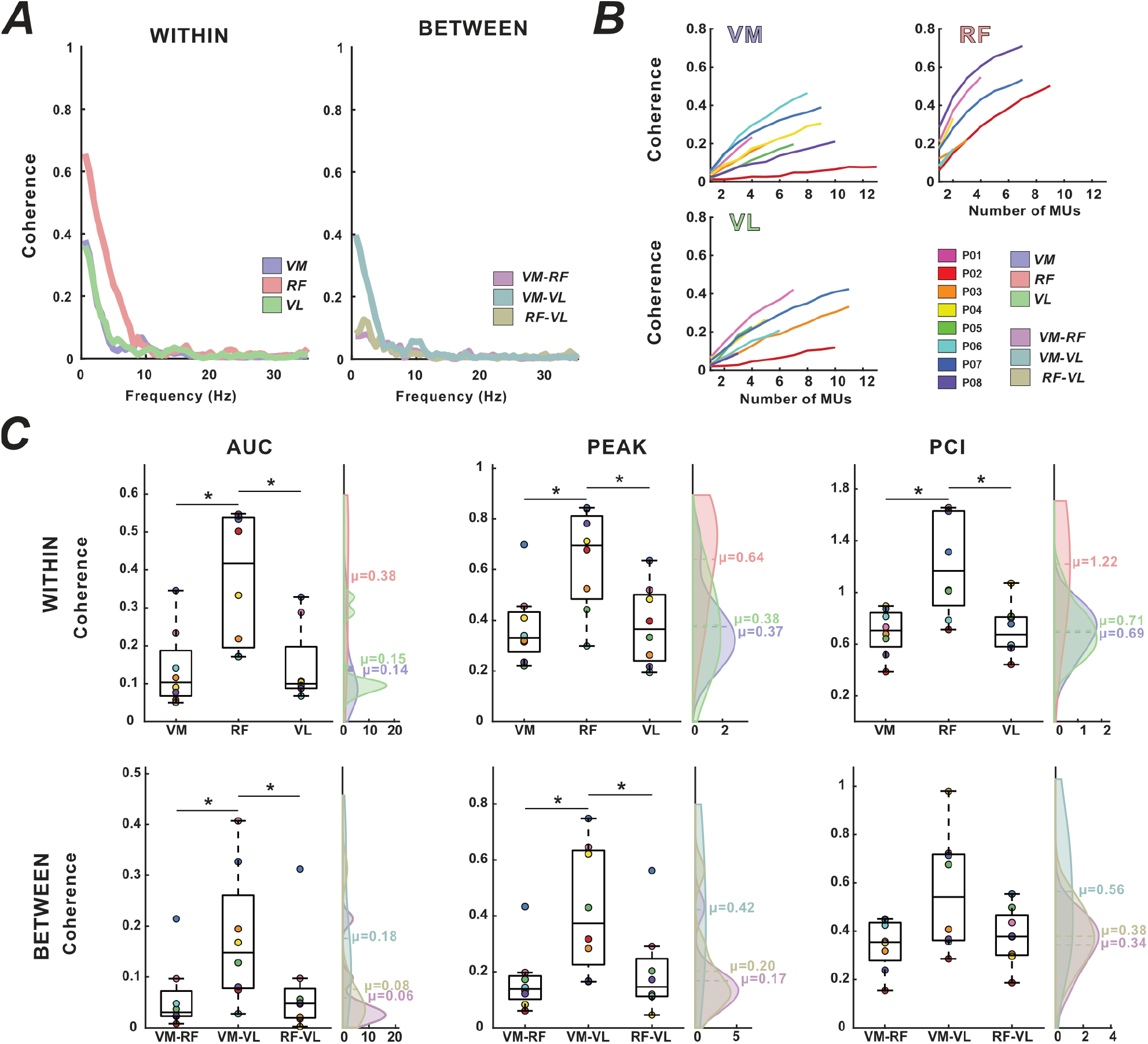
(A) Coherence was estimated between two equally sized subpools of motor units (MUs), each summed into a cumulative spike train (CST). This procedure was repeated 100 times with random subpool assignments and the resulting coherence estimates averaged. Within-muscle coherence was computed between two subpools from the same muscle (A, left panel); across-muscle coherence between subpools from different muscles (A, right panel). To ensure comparability across muscles, the number of MUs included per participant was limited to the lowest MU count across all three muscles. (B) Normalized area under the curve (AUC) as a function of increasing pool size for within-muscle conditions in the delta band (0–5 Hz). Different colors represent different participants. (C) Summary coherence metrics AUC, peak coherence, and percentage of common input (PCI) for the delta band for within-muscle (upper row) and across-muscle (lower row), displayed as box plots with overlaid data points and a marginal density plot. Asterisks (*) denote statistically significant differences following Holm correction (*α* = 0.05).

### Cumulative Explained Variance

PCA and FA were applied to the Hanning-convolved, z-scored spike trains to characterize the dimensionality of each MU pool. The first eight components or factors were retained, and MUs were subsampled to the minimum MU count to ensure comparability across muscles. For PCA, a significant main effect of muscle was observed, *F*(2, 186) = 9.4, *p <* 0.001, Cohen’s *f* ^2^ = 0.10, as well as a significant muscle *×* component interaction, *F*(2, 186) = 3.24, *p* = 0.041, Cohen’s *f* ^2^ = 0.03, indicating that the rate at which cumulative variance accumulated differed between muscles (Figure 5 PCA). Post-hoc comparisons revealed that RF required fewer components to explain the same proportion of variance compared to both VM (components 1 to 5, all *p <* 0.05, *d* = 0.6 to 1.3) and VL (components 1 to 5, all *p <* 0.05, *d* = 0.6 to 1.3), whereas VM and VL did not differ at any component level. FA showed a similar overall pattern. A significant main effect of muscle was observed, *F*(2, 186) = 36.7, *p <* 0.001, Cohen’s *f* ^2^ = 0.40, whereas the × muscle factor interaction was not significant, *F*(2, 186) = 0.72, *p* = 0.489, Cohen’s *f* ^2^ = 0.01, indicating that the difference in cumulative variance between muscles was stable across the number of factors retained. Post-hoc comparisons revealed that RF required fewer factors to explain the same proportion of variance compared to both VM (factors 1 to 8, all *p <* 0.05, *d* = 2.2 to 2.6) and VL (factors 1 to 8, all *p <* 0.05, *d* = 2.3 to 3.0), whereas VM and VL did not differ at any factor level. Together, both approaches converged on the same pattern. RF required fewer components or factors than VM or VL to reach comparable levels of explained variance.

**Figure 5.**
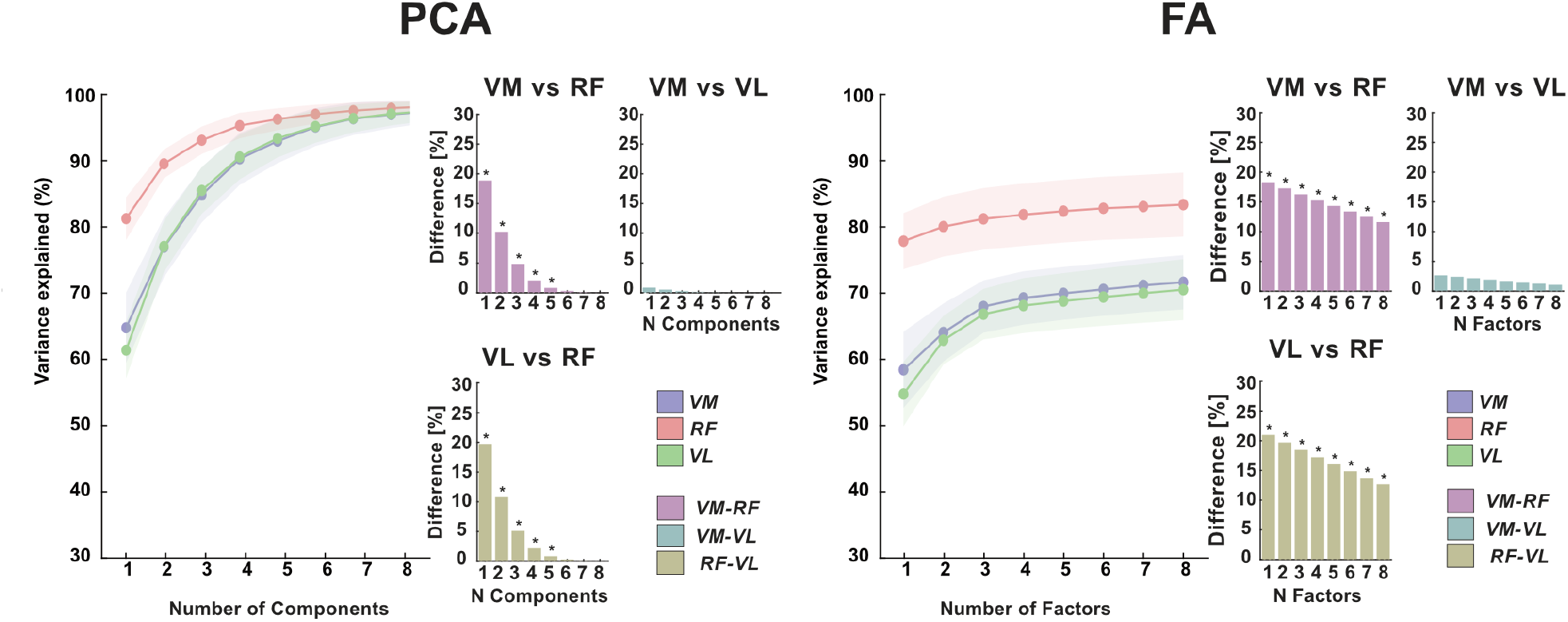
Cumulative explained variance as a function of the number of principal components (PCA, left) or latent factors (FA, right) retained, shown for each muscle (VM, RF, VL) and each pairwise muscle comparison (VM vs. RF, VM vs. VL, VL vs. RF). Curves with shaded areas (mean ± SEM) represent the cumulative variance explained for each muscle. Bars indicate the difference in cumulative explained variance between the two muscles of each pair at the respective component or factor level. Asterisks (*) indicate significant pairwise differences following Holm correction (*α* = 0.05).

### Factor Analysis

Factor analysis was applied to the pooled spike trains of all MUs across VM, RF and VL collectively (Figure 6 A). The optimal number of latent factors was determined individually for each participant, after which each factor was assigned to a dominant muscle based on its loading profile, and individual MUs were assigned to the muscle factor on which they showed the highest loading (Figure 6 D). The optimal number of factors varied across participants, ranging from 2 to 4 (Figure 6 B). When only two factors were extracted, VM and VL MUs were jointly represented by a single shared factor. In solutions with three or more factors, VM and VL MUs were distributed across multiple factors with varying proportions, rather than being exclusively represented by a single muscle-specific factor. On average, the entire RF MU pool was exclusively explained by the RF-factor (100 ± 0%), with no contribution from the VM- or VL-factor. Within the VM MU pool, 89.9 ± 11.2% of MUs were explained by the VM-factor and 8.0 ± 10.0% by the VL-factor, with absent contribution from the RF-factor (0%). Conversely, within the VL MU pool, 59.5 ± 22.7% of MUs were explained by the VL-factor and 25.3 ± 24.0% by the VM-factor, again with absent contribution from the RF-factor (0%). The percentage of MUs dominated by each muscle-specific factor is illustrated in Figure 6 C.

**Figure 6.**
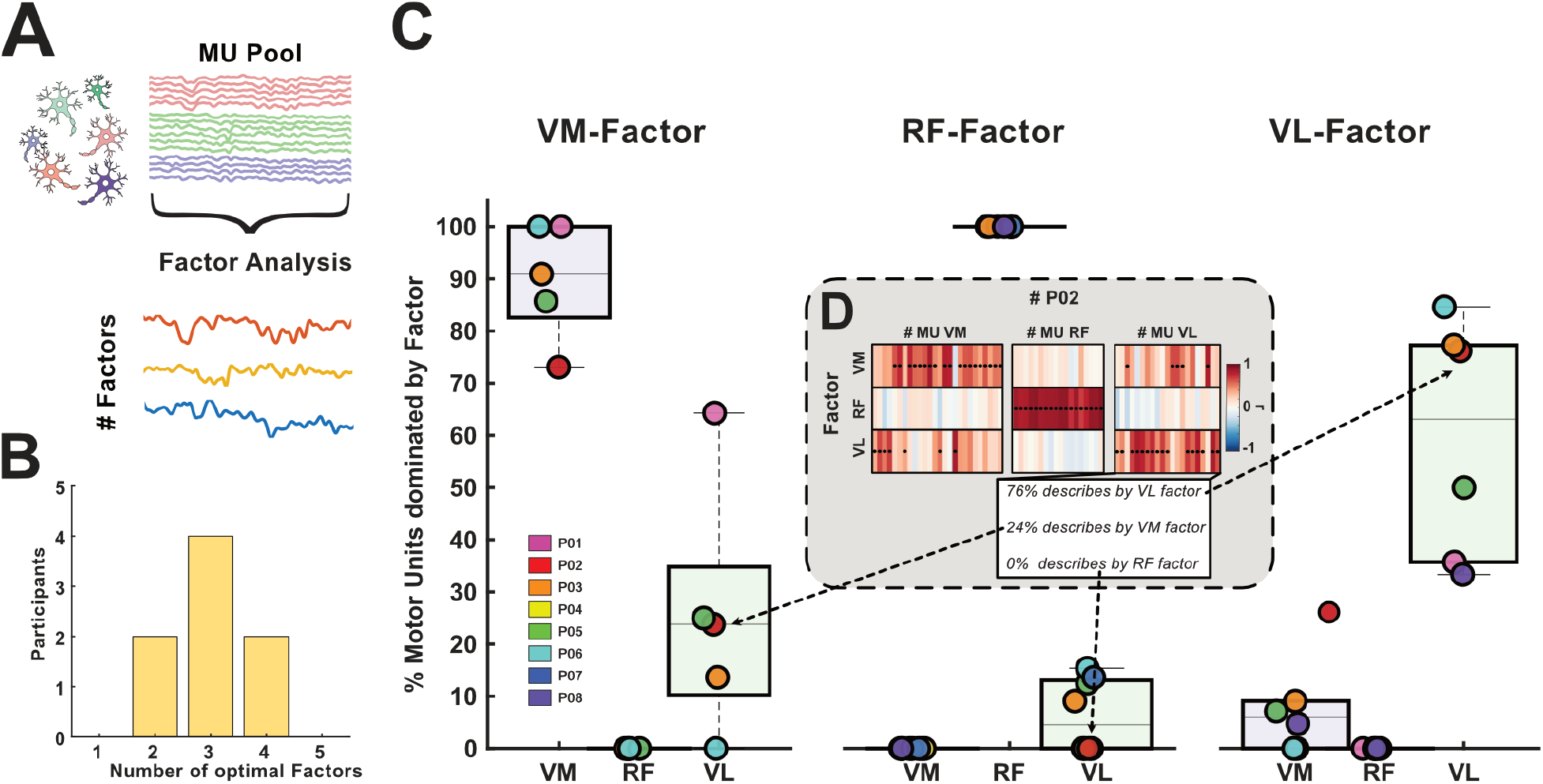
(A) Factor analysis was applied to the pooled motor unit (MU) spike trains across all three muscles (VM, RF, VL) to identify latent factors best describing the combined MU pool. (B) The optimal number of factors varied across participants, with the majority best described by three factors. (C) Percentage of MUs from each muscle (VM, RF, VL) assigned to the muscle-specific factor (VM–Factor, RF–Factor, VL–Factor), shown as box plots with individual participant indicated by different colors. The MUs of RF were exclusively explained by the RF–Factor, whereas the VM–Factor and VL–Factor each predominantly captured MUs of their respective muscle but also contained a notable proportion of MUs from the other vastii muscle, reflecting the more variable and overlapping input structure between VM and VL. (D) Representative example for one participant (P 02) illustrating how the percentages in (C) were derived: following factor-to-muscle assignment, each MU was correlated with every extracted factor, and the percentage of MUs predominantly explained by each factor was calculated per muscle.

## DISCUSSION

The present study characterized the neural control of three quadriceps muscles, VM, RF, and VL during isometric knee extension at 30% MVC, using multiple intramuscular EMG recordings. Several complementary metrics, including discharge characteristics, pairwise correlation, spectral coherence, explained variance, and factor analysis, were applied to assess the organization of common neural input within and across muscles. Despite their shared mechanical output via a common patellar insertion, the results consistently reveal that RF receives a strong, homogeneous, common input that is largely independent of the vasti, while VM and VL share a variable degree of common drive whose organization differs between individuals.

### Discharge Characteristics

A recent systematic review and meta-analysis on MU discharge behavior across a range of contraction intensities reported generally lower DR for most lower-limb muscles compared with muscles of the upper extremity (Inglis et al., 2025). Although RF was mentioned in the review, it was excluded from quantitative synthesis due to insufficient data of only two studies. The RF DR observed here exceeds those of VM and VL and are more comparable to values reported for upper-limb muscles, challenging a straightforward limb-based categorization. This pattern is more consistent with functional demands than with limb membership. Muscles involved in fine motor control, such as the FDI (∼14 Hz) and biceps brachii (∼14 Hz), require higher DR to achieve greater twitch fusion (Milner-Brown et al., 1973; Nordstrom et al., 1989), whereas muscles with primarily postural roles, such as the soleus (SOL) (∼9 Hz) and the vasti (∼10 Hz), rely more on recruitment than on rate modulation (De Luca et al., 1982; Honeine et al., 2013; Vieira et al., 2012). The higher CoV observed in RF is consistent with a stronger rate coding contribution (Negro et al., 2009; Farina and Negro, 2015). Although direct comparisons are limited by sparse data, the available literature on other muscles with dynamic force-generating roles, such as biceps femoris and semitendinosus (Kirk and Rice, 2017; Kirk et al., 2018; Sahinis et al., 2025), similarly reports elevated discharge rates, pointing in the same direction.

### Independent and Homogeneous Neural Drive to the Rectus Femoris

The converging evidence from pairwise correlation, spectral coherence, explained variance, and factor analysis consistently characterizes the RF MN pool as being governed by a single common input that is largely independent of the drive to the vasti. Correlation values were markedly higher within RF than within either vasti muscle, whereas correlations involving RF remained low between muscles. In contrast, within- and between-muscle correlations for VM and VL were similar, with only a slight reduction in the between-muscle correlations. Due to the input–output nonlinearity of individual MN, the strength of common synaptic input is not proportional to the magnitude of correlation between pairs of discharge times (Farina and Holobar, 2016), and the absence of significant pairwise correlation within VM and VL can therefore not be taken as evidence against the presence of common input to those pools (Negro and Farina, 2012). The consistently elevated within-RF correlations across all recorded units and all participants do, however, provide converging evidence for a strong and uniformly distributed common input to the RF pool.

Spectral coherence confirmed the correlation results in the frequency domain (Figure 4). RF exhibited high within-muscle but low between-muscle coherence, whereas VM and VL showed similar within- and between-muscle coherence, consistent with the correlation analysis and previous observations (Rossato et al., 2022). It should be noted that coherence estimates scale with pool size (Negro and Farina, 2012; Farina et al., 2014b; Dideriksen et al., 2018; Castronovo et al., 2015) and the effective pool size differed considerably across participants in the present study. PCI values for RF exceeded 1 in several participants, likely reflecting a steep initial rise in the coherence-versus-unit-count curve with insufficient units to reach saturation, resulting in an unreliable fit. While this represents a limitation of the PCI metric when applied to small MU pools, the consistently elevated PCI values, some exceeding 1.6, reinforce the interpretation of strong common input to RF, as such values arise precisely because the coherence curve rises steeply even with few units.

PCA and FA provide complementary perspectives on the dimensionality of common synaptic input. While PCA explains all variance once the number of components equals the number of MU, FA partitions the variance into shared and unique components and therefore quantifies only the common variance. In both approaches, RF exhibited a higher explained variance with a single component/factor than VM or VL. This difference was particularly evident in the FA, where the explained variance for VM and VL increased substantially with the first three factors before reaching a plateau, whereas RF saturated after the first factor with only minor gains from additional factors. Consistent with this observation, factor loading analysis assigned all RF MUs exclusively to an RF-specific factor across all participants and factor solutions, and t-SNE further visualized RF as a distinct cluster in the low-dimensional representation.

That RF can be activated independently of VM and VL was previously suggested indirectly at the whole-muscle level using surface EMG under experimental pain (Hug et al., 2014), and its functional independence is therefore not unexpected given its biarticular anatomy and additional role in hip flexion. However, differences in muscle anatomy alone do not necessarily imply completely segregated neural inputs. Using PCA, Levine et al. (2023) showed that the monoarticular SOL receives shared input from the biarticular gastrocnemii, while GM and GL do not share common input. The present study extends this to the MN level and demonstrates that, unlike the SOL, the entire RF MN pool is governed by a single, homogeneous common input that is segregated from those of VM and VL during isometric leg extensions. Recent studies further suggest that the organization of common synaptic input is even more spatially refined than previously assumed. Weinman et al. (2026) identified separate input components for proximal and distal regions of TA using HDsEMG, while Haller et al. (2026) demonstrated with targeted intramuscular recordings that MU in proximal VM shared greater similarity with distal VL than with distal VM. These findings indicate that muscle architecture and regional function, rather than muscle identity alone, shape the organization of common synaptic input. Importantly, the present recordings sampled MU across the entire proximodistal extent of each quadriceps muscle using multiple targeted intramuscular electrodes. Despite capturing this regional variability, RF consistently remained a distinct MN population across all analyses.

### Individual Differences within and across Muscles

Despite RF consistently showing the highest within-muscle correlation and coherence, substantial inter-individual variability was present. Comparing correlation structures across individuals revealed that some participants exhibited an elevated overall correlation level and a steeper increase in coherence with increasing pool size than others, indicating that the underlying neural strategies differ between individuals. Crouzier et al. (2019) documented stable inter-individual variability in quadriceps and triceps surae activation distributions across sessions and tasks, concluding that this variability reflects genuine neural strategies rather than measurement noise. Avrillon et al. (2021) extended this to the MU level, demonstrating that the proportion of common drive between VM and VL, as well as the degree of muscle-specific drive, varied substantially across individuals, with values ranging from near-complete sharing to near-complete independence. Crucially, this variability was not associated with training level or sport participation (Crouzier et al., 2019), leaving its origin unresolved. Fiber-type composition is a further candidate, as slow- and fast-twitch MU may differ in their susceptibility to common synaptic input, but no validated method currently exists to classify MU fiber-type from discharge patterns alone (Heckman and Enoka, 2012), leaving open whether differences in common input organization reflect the differential recruitment of distinct fiber-type populations.

Spinal inhibitory circuits represent another plausible mechanism. Using paired-pulse H-reflex conditioning of the SOL, Matsuya et al. (2017) demonstrated that the strength of recurrent inhibition correlated positively with corticomuscular coherence across individuals, arguing that recurrent inhibition via Renshaw cells is a source of interindividual variation in oscillatory neural drive. Whether an analogous relationship holds for the quadriceps remains unknown. More recently, Hug et al. (2025) demonstrated that experimental knee pain reduced the proportion of common synaptic input to VL motor units and provided evidence for a non-homogeneous distribution of inhibitory inputs across the motor neuron pool. Further, Dernoncourt et al. (2025) showed that heteronymous Renshaw cell connectivity between MN pools can increase the dimensionality of population activity even under a single descending command, but only when pools already share substantial common input. The chronic mechanical co-activation of VM and VL may therefore promote stronger heteronymous recurrent inhibition between these pools than between RF and the vasti, and inter-individual differences in the strength of this connectivity could directly account for the variable factor structures observed.

### Limitations

Several limitations should be noted. First, the final sample comprised only eight participants after excluding two individuals with insufficient RF MU, limiting statistical power and generalization of the observed inter-individual variability. Second, RF pools contained fewer MUs than VM and VL, although coherence and variance measures were matched for pool size, smaller pools reduce estimation precision and likely contributed to the PCI values exceeding 1 observed in several participants. Third, signal quality varied across the implanted wire electrodes. The inter-wire distance of each bipolar pair can shift during implantation and with muscle contraction, which occasionally reduced the signal-to-noise ratio and hampered MU decomposition at specific sites. As the affected recording sites (proximal, central, or distal) differed across participants, we refrained from performing systematic within-muscle comparisons between regions and instead analyzed pooled MU data per muscle. Fourth, correlation, coherence, explained variance, and factor analysis characterize the statistical structure of shared input but cannot identify its physiological origin (e.g., descending drive vs. spinal circuitry). Finally, recordings were limited to an isometric task at a fixed force level and joint angle, and results may not extend to other contraction tasks.

### SUMMARY

The present study provides a multi-method characterization of the neural input organization across three quadriceps muscles during isometric knee extension. Despite their shared mechanical function as a knee extensor, VM, RF, and VL were governed by distinct neural strategies. RF was characterized by higher discharge rates, stronger and more low-dimensional common input, and an invariant dedicated latent factor structure across all participants, likely reflecting its functional specialization as a biarticular muscle. To the best of our knowledge, this is the first study to directly demonstrate a fully independent input to a muscle classically considered synergistic with the vastii muscles. VM and VL, by contrast, shared a variable degree of common drive with factor solutions ranging from a single shared factor to largely independent muscle-specific inputs. Together, these findings highlight that neural input architecture within a synergist group reflects functional specialization rather than mechanical coupling alone, and underscore the need for individual-level rather than group-averaged analysis when characterizing MN pool organization.

## GRANT

This work was supported by the European Research Council (ERC) through the project GRASPAGAIN under grant 101118089, and by the German Research Foundation (DFG) through the project DeMOTUS under grant 523352235 to A.D V..

## AUTHOR CONTRIBUTIONS

F.B., D.H., and A.D.V. conceived and designed research; F.B., D.H., L.H., S.Z. performed experiments; F.B., and A.D.V.analyzed data; F.B., D.H., L.H., S.Z., M.B., and A.D.V. interpreted results of experiments; F.B., and A.D.V. prepared figures; F.B., and A.D V. drafted manuscript; F.B., D.H., L.H., S.Z., M.B., and A.D.V. edited and revised manuscript; F.B., D.H., L.H., S.Z., M.B. and A.D.V. approved final version of manuscript.

## REFERENCES

Avrillon, S., Del Vecchio, A., Farina, D., Pons, J. L., Vogel, C., Umehara, J., and Hug, F. (2021). Individual differences in the neural strategies to control the lateral and medial head of the quadriceps during a mechanically constrained task. Journal of Applied Physiology, 130:269–281.

Bräcklein, M., Barsakcioglu, D. Y., Ibáñez, J., Eden, J., Burdet, E., Mehring, C., and Farina, D. (2022). The control and training of single motor units in isometric tasks are constrained by a common input signal. eLife, 11:e72871.

Buja, A. and Eyuboglu, N. (1992). Remarks on parallel analysis. Multivariate Behavioral Research, 27(4):509–540.

Cabral, H. V., Cudicio, A., Bonardi, A., Del Vecchio, A., Falciati, L., Orizio, C., Martinez-Valdes, E., and Negro, F. (2024). Neural filtering of physiological tremor oscillations to spinal motor neurons mediates short-term acquisition of a skill learning task. eNeuro, 11(7):ENEURO.0043–24.2024.

Castronovo, A. M., Negro, F., Conforto, S., and Farina, D. (2015). The proportion of common synaptic input to motor neurons increases with an increase in net excitatory input. Journal of Applied Physiology, 119:1337–1346.

Clark, D. J., Ting, L. H., Zajac, F. E., Neptune, R. R., and Kautz, S. A. (2010). Merging of healthy motor modules predicts reduced locomotor performance and muscle coordination complexity post-stroke. Journal of Neurophysiology, 103(2):844–857.

Crouzier, M., Hug, F., Dorel, S., Deschamps, T., Tucker, K., and Lacourpaille, L. (2019). Do individual differences in the distribution of activation between synergist muscles reflect individual strategies? Experimental Brain Research, 237:625–635.

De Luca, C. J. and Erim, Z. (1994). Common drive of motor units in regulation of muscle force. Trends in Neurosciences, 17(7):299–305.

De Luca, C. J., LeFever, R. S., McCue, M. P., and Xenakis, A. P. (1982). Control scheme governing concurrently active human motor units during voluntary contractions. The Journal of Physiology, 329:129–142.

Del Vecchio, A., Marconi Germer, C., Kinfe, T. M., Nuccio, S., Hug, F., Eskofier, B., Farina, D., and Enoka, R. M. (2023). The forces generated by agonist muscles during isometric contractions arise from motor unit synergies. The Journal of Neuroscience, 43:2860–2873.

Dernoncourt, F., Avrillon, S., Logtens, T., Cattagni, T., Farina, D., and Hug, F. (2025). Flexible control of motor units: is the multidimensionality of motor unit manifolds a sufficient condition? The Journal of Physiology, 603(8):2349–2368.

Dideriksen, J. L., Negro, F., Falla, D., Kristensen, S. R., Mrachacz-Kersting, N., and Farina, D. (2018). Coherence of the surface emg and common synaptic input to motor neurons. Frontiers in Human Neuroscience, 12:207.

Farina, D. and Holobar, A. (2016). Characterization of Human Motor Units From Surface EMG Decomposition. Proceedings of the IEEE, 104(2):353–373.

Farina, D., Merletti, R., and Enoka, R. M. (2014a). The extraction of neural strategies from the surface emg: an update. Journal of Applied Physiology, 117:1215–1230.

Farina, D. and Negro, F. (2015). Common synaptic input to motor neurons, motor unit synchronization, and force control. Exercise and Sport Sciences Reviews, 43:23–33.

Farina, D., Negro, F., and Dideriksen, J. L. (2014b). The effective neural drive to muscles is the common synaptic input to motor neurons. The Journal of Physiology, 592:3427–3441.

Gandevia, S. C. (2001). Spinal and supraspinal factors in human muscle fatigue. Physiological Reviews, 81(4):1725– 1789.

Germer, C. M., Farina, D., Elias, L. A., Nuccio, S., Hug, F., and Vecchio, A. D. (2021). Surface emg cross talk quantified at the motor unit population level for muscles of the hand, thigh, and calf. Journal of Applied Physiology, 131:808–820.

Haller, D., Beermann, F., Sîmpetru, R. C., Hofbeck, L., Enoka, R. M., and Del Vecchio, A. (2026). Voluntary dissociation of motor unit activity in the vastii muscles. The Journal of Neuroscience, 46(21):e1982252026.

Heckman, C. J. and Enoka, R. M. (2012). Motor unit. Comprehensive Physiology, 2:2629–2682.

Holobar, A. and Zazula, D. (2007). Multichannel blind source separation using convolution kernel compensation. IEEE Transactions on Signal Processing, 55(9):4487–4496.

Honeine, J.-L., Schieppati, M., Gagey, O., and Do, M.-C. (2013). The functional role of the triceps surae muscle during human locomotion. PLoS ONE, 8:e52943.

Horn, J. L. (1965). A rationale and test for the number of factors in factor analysis. Psychometrika, 30(2):179–185.

Hug, F., Avrillon, S., Sarcher, A., Del Vecchio, A., and Farina, D. (2023). Correlation networks of spinal motor neurons that innervate lower limb muscles during a multi-joint isometric task. The Journal of Physiology, 601:3201–3219.

Hug, F., Del Vecchio, A., Avrillon, S., Farina, D., and Tucker, K. (2021). Muscles from the same muscle group do not necessarily share common drive: evidence from the human triceps surae. Journal of Applied Physiology, 130:342–354.

Hug, F., Dernoncourt, F., Avrillon, S., Thorstensen, J., Besomi, M., Van Den Hoorn, W., and Tucker, K. (2025). Non-homogeneous distribution of inhibitory inputs among motor units in response to nociceptive stimulation at moderate contraction intensity. The Journal of Physiology, 603:3445–3461.

Hug, F., Hodges, P. W., Hoorn, W. V. D., and Tucker, K. (2014). Between-muscle differences in the adaptation to experimental pain. Journal of Applied Physiology, pages 1132–1140.

Inglis, J. G., Cabral, H. V., Cosentino, C., Bonardi, A., and Negro, F. (2025). Motor unit discharge behavior in human muscles throughout force gradation: a systematic review and meta-analysis with meta-regression. Journal of Applied Physiology, 138:1050–1065.

Kirk, E. A., Gilmore, K. J., and Rice, C. L. (2018). Neuromuscular changes of the aged human hamstrings. Journal of Neurophysiology, 120(2):480–488.

Kirk, E. A. and Rice, C. L. (2017). Contractile function and motor unit firing rates of the human hamstrings. Journal of Neurophysiology, 117:243–250.

Laine, C. M., Martinez-Valdes, E., Falla, D., Mayer, F., and Farina, D. (2015). Motor neuron pools of synergistic thigh muscles share most of their synaptic input. The Journal of Neuroscience, 35:12207–12216.

Lee, M.-J., Ofner, P., Huang, H.-Y., Mulder, J., Ibañez, J., Farina, D., and Mehring, C. (2026). Rigid control of motor unit firing rates in the human tibialis anterior muscle persists during neurofeedback. Journal of Neurophysiology, 135:1499–1517.

Levine, J., Avrillon, S., Farina, D., Hug, F., and Pons, J. L. (2023). Two motor neuron synergies, invariant across ankle joint angles, activate the triceps surae during plantarflexion. The Journal of Physiology, 601:4337–4354.

Liddell, E. G. T. and Sherrington, C. S. (1925). Recruitment and some other features of reflex inhibition. Proceedings of the Royal Society of London. Series B, Containing Papers of a Biological Character, 97(686):488–518.

Maillet, J., Avrillon, S., Nordez, A., Rossi, J., and Hug, F. (2022). Handedness is associated with less common input to spinal motor neurons innervating different hand muscles. Journal of Neurophysiology, 128:778–789.

Matsuya, R., Ushiyama, J., and Ushiba, J. (2017). Inhibitory interneuron circuits at cortical and spinal levels are associated with individual differences in corticomuscular coherence during isometric voluntary contraction. Scientific Reports, 7:44417.

McGill, K. C., Lateva, Z. C., and Marateb, H. R. (2005). EMGLAB: an interactive EMG decomposition program. Journal of Neuroscience Methods, 149:121–133.

Milner-Brown, H. S., Stein, R. B., and Yemm, R. (1973). Changes in firing rate of human motor units during linearly changing voluntary contractions. The Journal of Physiology, 230:371–390.

Myers, L. J., Erim, Z., and Lowery, M. M. (2004). Time and frequency domain methods for quantifying common modulation of motor unit firing patterns. Journal of NeuroEngineering and Rehabilitation, 1:2.

Negro, F. and Farina, D. (2012). Factors influencing the estimates of correlation between motor unit activities in humans. PLoS ONE, 7(9):e44894.

Negro, F., Holobar, A., and Farina, D. (2009). Fluctuations in isometric muscle force can be described by one linear projection of low-frequency components of motor unit discharge rates. The Journal of Physiology, 587:5925–5938.

Negro, F., Muceli, S., Castronovo, A. M., Holobar, A., and Farina, D. (2016a). Multi-channel intramuscular and surface EMG decomposition by convolutive blind source separation. Journal of Neural Engineering, 13(2):026027.

Negro, F., Yavuz, U., and Farina, D. (2016b). The human motor neuron pools receive a dominant slow-varying common synaptic input. The Journal of Physiology, 594(19):5491–5505.

Nordstrom, M. A., Miles, T. S., and Veale, J. L. (1989). Effect of motor unit firing pattern on twitches obtained by spike-triggered averaging. Muscle Nerve, 12:556–567.

Rossato, J., Avrillon, S., Tucker, K., Farina, D., and Hug, F. (2024). The volitional control of individual motor units is constrained within low-dimensional neural manifolds by common inputs. The Journal of Neuroscience, 44(34):e0702242024.

Rossato, J., Tucker, K., Avrillon, S., Lacourpaille, L., Holobar, A., and Hug, F. (2022). Less common synaptic input between muscles from the same group allows for more flexible coordination strategies during a fatiguing task. Journal of Neurophysiology, 127:421–433.

Sahinis, C., Amiridis, I. G., Farina, D., Enoka, R. M., and Kellis, E. (2025). Independent neural drives and distinct motor unit discharge characteristics in hamstring muscles during isometric knee flexion. European Journal of Applied Physiology, 126(2):839–852.

van der Maaten, L. and Hinton, G. (2008). Visualizing data using t-sne. Journal of Machine Learning Research, 9(86):2579–2605.

Vieira, T. M. M., Loram, I. D., Muceli, S., Merletti, R., and Farina, D. (2012). Recruitment of motor units in the medial gastrocnemius muscle during human quiet standing: is recruitment intermittent? what triggers recruitment? Journal of Neurophysiology, 107:666–676.

Weinman, L. E., Del Vecchio, A., Mazzo, M. R., and Enoka, R. M. (2024). Motor unit modes in the calf muscles during a submaximal isometric contraction are changed by brief stretches. The Journal of Physiology, 602:1385–1404.

Weinman, L. E., Lykidis, A., Amiridis, I. G., Sahinis, C., and Enoka, R. M. (2026). Two anatomically distinct motor unit modes in tibialis anterior during submaximal isometric contractions. Journal of Neurophysiology, 135:214–226.

